# Effects of Different Microplastics on Nematodes in the Soil Environment: Tracking the Extractable Additives using an Ecotoxicological Approach

**DOI:** 10.1101/2020.07.07.192278

**Authors:** Shin Woong Kim, Walter R. Waldman, Matthias C. Rillig

**Affiliations:** Institute of Biology, Freie Universität Berlin, 14195 Berlin, Germany; Berlin-Brandenburg Institute of Advanced Biodiversity Research. 14195 Berlin, Germany; Department of Environmental Health Science, Konkuk University, 120 Neungdong-ro, Gwangjin-gu, Seoul 05029, Korea; Science and Technology Center for Sustainability, Federal University of São Carlos, 18052-780, Sorocaba/SP, Brazil

**Keywords:** *Caenorhabditis elegans*, Composition, Shape, Size, Toxicity

## Abstract

With an increasing interest in the effects of microplastic in the soil environment, there is a need to thoroughly evaluate potential adverse effects of these particles as a function of their characteristics (size, shape, and composition). In addition, extractable chemical additives from microplastic have been identified as an important toxicity pathway in the aquatic environment. However, we currently know little about effects of such additives in the soil environment. In this study on nematodes (*Caenorhabditis elegans*), we adopted an ecotoxicological approach to assess the potential effects of thirteen different microplastics with different characteristics and extractable additives. We found that toxic effects appear to increase in the order of low-density polyethylene (LDPE) film < polypropylene (PP) fragments < high-density polyethylene (HDPE) fragments ≈ polystyrene (PS) fragments < polyethylene terephthalate (PET) fragments ≈ polyacrylicnitrile (PAN) fibers. Acute toxicity was mainly attributed to the extractable additives: when the additives were extracted, the toxic effects of each microplastic disappeared in the acute soil toxicity test. The harmful effects of LDPE film and PAN fibers increased when the microplastics were maintained in soil for a long-term period with frequent wet-dry cycles. We here provide clear evidence that microplastic toxicity in the soil is highly related to particle characteristics and extractable additives. Our results suggest that future experiments consider extractable additives as a key explanatory variable.

**Abstract art/Table of contents:** 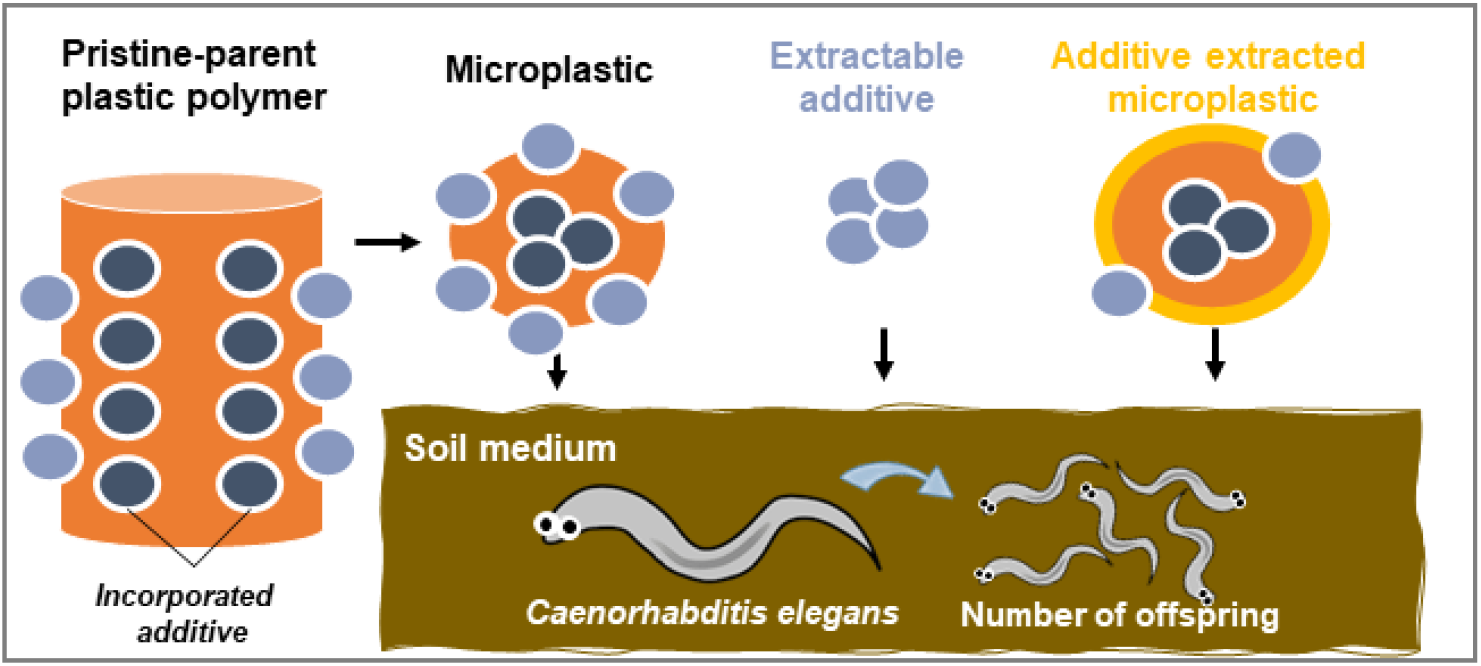

## INTRODUCTION

Plastic polymers have been widely used during the last 70 years, and an enormous amount of plastic litter has been spread into the environment.^1,2^ Primary plastics have been fragmented into a smaller size (<5 mm, microplastic), and these tiny particles are ubiquitously detected in a broad range of environmental compartments including ocean,^1^ freshwater,^3^ soil,^4^ atmosphere,^5^ and even our drinking water.^6^ Although microplastic pollution in the soil environment has not received a similar media and research attention, microplastic abundances are estimated to be up to 23-times larger than those in the ocean.^7^ Soils have various input sources including amendment and irrigation with microplastics,^8^ and previous studies have reported that 8 to 67,500 mg kg^−1^ of microplastics can be observed at industrial,^9^ pristine floodplain,^10^ and agricultural lands.^11^

A key concern of microplastic pollution is whether it poses a risk to ecosystems. Although lack of available data and methodological issues still limit progress,^12^ several previous studies have provided laboratory-scale evidence of harmful effects on living organisms.^4,13^ In the soil environment, invertebrates and agricultural plants can experience adverse effects, such as mortality and growth decrease,^14–17^ and negative effects on microbial and enzyme activities have also been reported.^18,19^ With an increasing number of these studies, it is becoming necessary to systematically test for what parameters control these effects,^4^ and microplastic characteristics (size, shape and composition) have been highlighted as an important factor to consider.^20,21^ While the database on microplastic effects is much stronger for aquatic environments,^22–25^ fewer such studies have been reported for soil. Several studies have reported size or composition-dependent effects on plant, nematode, and soil properties,^14,26–31^ but a part of these studies were performed in non-soil media or using spherical beads.^26,27,29,30^

Chemical effects may serve as an important mediator of microplastic toxicity. A central hypothesis is that microplastic can carry harmful hydrophobic organic pollutants with a strong sorption capacity,^32^ and that leaching of chemical additives from microplastic can be expected.^33^ These additives are intentionally added to plastic products to improve their functionality (*e.g.*, functional additives, colorants, fillers, and reinforcements), and are optimized for the first use phase, not for recycling.^34,35^ These incorporated chemicals can be continually released into the environment during the decomposition or fragmentation process, and might be partially responsible for any microplastic toxicity.^34,36^ For example, bisphenol A, which is authorized under Registration, Evaluation, Authorisation & Restriction of Chemicals (REACH) as a stabilizer, is regarded as an estrogen agonist.^35^ Phthalates are another common organic species in plastic manufacturing,^37^ and they are considered endocrine disruptors at very low environmental levels (in the range of ng L^−1^).^38^ Several experiments have found that leaching solutions from microplastic can induce severe damage on aquatic organisms including water fleas,^39,40^ microalgae,^41,42^ copepods,^43^ and brown mussels,^44^ while there are no data reporting harmful effects of plastic additives in the soil environment.

Since additive leaching from plastics is highly related to both chemical equilibria and diffusion kinetics, a partition constant (K_D_) between plastic and surrounding media can be the most important factor to understand the leaching mechanism.^32,33^ Nevertheless, K_D_ is mostly calculated with pure solvents or food simulants and non-degraded polymers, having limited information of K_D_ for microplastic research, considering secondary microplastics or environmental conditions.^32,45,46^ Furthermore, the immediate surrounding of microplastics in aquatic environments is dynamic, constantly changing due to physicochemical and biological parameters.^47,48^ Several pieces of evidence have also suggested that microplastics might be transported from the surface into the soil system through cracking or movement of living organisms,^49,50^ and the physicochemical properties of surrounding media are varied and complicated similar to aquatic environments.^51–54^ Since we have no knoweldge to predict the effects of chemical additive in such soil media from first principles, data from experimental studies are needed.

Here, we conducted soil toxicity tests using the nematode *Caenorhabditis elegans* as a model organism, and thirteen microplastics were chosen as target materials; six different compositions (high-density polyethylene, HDPE; polyethylene terephthalate, PET; polypropylene, PP; polystyrene, PS; low-density polyethylene, LDPE; polyacrylicnitrile, PAN) and three different shapes (fragments; film; fibers) with one to three different size ranges (Table S1). In order to evaluate potential effects of extractable additives from each microplastic, we adopted an ecotoxicological approach instead of prediction by chemical analysis. The additives were extracted with water as solvent using two different methods. The most efficient method was used to follow the influence of microplastics size and concentration in the ecotoxicological assessment. Finally, microplastic ecotoxicity was tested in two situations: afte removing additives from partical surfaces, to correlate toxicity and the presence of the additive, and an ecotoxicological assessment as a function of time with soil experiencing wet-dry cycles.

## MATERIALS AND METHODS

### Target Microplastics and Organism

Target microplastic fragments were prepared by cryo-milling as reported in our previous study.^14^ The polymers, including HDPE, PET, PP, and PS were obtained from Bundesanstalt für Materialforschung und -prüfung (Berlin, Germany), and they were ground in an ultra-centrifugal mill after embrittlement with liquid nitrogen. After drying, the fragments were passed through a 1000 μm-sieve and stored at room temperature. In order to obtain different sized fragments, sieving (630 and 250 μm) was additionally performed in the present study. HDPE, PP, and PS were prepared with three different size ranges (<250, 250-630, and 630-1000 μm), and PET was separated into two size ranges (<250 and 250-630 μm) due to smaller size distribution than others.^14^ LDPE film and PAN fibers were prepared using commercial mulching film (LDPE; thickness, 13.66 ± 2.32 μm, Ihlshin Chemical Co., Ltd., Ansan, South Korea) and knitting wool (100% PAN, DIKTAS Sewing & Knitting Yarns Co., Turkey), respectively.^55^ Each material was cut using sterilized scissors, and then passed through a 630 μm-sieve (<630 μm) and stored at room temperature. For the spectroscopical characterization we used a spectrophotometer (Jasco, model FT/IR-4100, ATR mode). Each sample was scanned 32 times, from 4000 to 600 cm^−1^, with resolution of 4 cm^−1^ (Figure S1).

*C. elegans* (wild type, Bristol strain N2) was obtained from Berlin Institute for Medical Systems Biology at the Max Delbrück Center for Molecular Medicine (Berlin, Germany). They were maintained on nematode growth medium (NGM; NaCl 3 g/L, peptone 2.5 g/L, agar 17 g/L, 1 M potassium phosphate 25 mL/L, 1 M CaCl_2_∙2H_2_O 1 mL/L, 1 M MgSO_4_∙7H_2_O 1 mL/L, cholesterol 1 mL/L) at 20 ± 2°C in the dark, and *Escherichia coli* (strain OP50) was supplied as a food source.^56^ In order to synchronize developmental stage before the experiment, the culture plates that were maintained for at least 3 days were treated with a Clorox solution (1 N NaOH:5% NaOCl, 1:1) for 20 min, and then the suspension containing embryos was centrifuged at 4500 rpm for 2 min. Subsequently, the embryo pellets were washed thrice with K-medium (0.032 M KCl, 0.051 M NaCl),^57^ and placed onto new NGM plate with *E. coli* strain OP50. The culture plates were incubated for 60-65 h for soil toxicity test.

### Soil Toxicity Tests

Test soil samples were collected from Linde, Märkisch Luch, Germany (52.545529N, 12.661135E) on April 18, 2018. The soil was passed through a 2 mm-sieve, and then dried at 60°C for 24 h. The texture of our test soil was a sand (sand 89.3%, silt 8.3%, and clay 2.4%), and pH and water holding capacity (WHC) were measured as 5.7 ± 0.2 and 0.32 ± 0.10 mL g^−1^, respectively (*n* = 3). In order to prepare test soils for microplastic fragments (HDPE, PET, PP, and PS) and film (LDPE), 100 mg of each microplastic was first mixed with 9.9 g of dry soil (1%), and then these initial mixtures were diluted using the same soil 10- and 100-times. Final test concentrations were determined as 0.01 (*n* = 4, fragments; *n* = 10, film), 0.1 (*n* = 4, fragments; *n* = 10, film), and 1 (*n* = 8, fragments; *n* = 10, film) % (based on dry weight in soil), and control sets (no microplastic added) were prepared with a matching equal number of replicates for every microplastic treatement set, respectively. For PAN fibers, 10 mg was mixed with 9.99 g of dry soil, and final test concentrations were 0.001 (*n* = 10), 0.01 (*n* = 10), and 0.1 (*n* = 10) %. Soil toxicity tests were performed as reported in previous studies.^28,58,59^ We added 0.3 g of microplastic-laced soil into each well of a 24-well plate, together with 76 μL of K-medium (80% of WHC). Ten age-synchronized worms were added to each well and maintained at 20 ± 2°C in the dark. After 24 h, soil containing nematodes was placed onto soil-agar isolation plates.^28,58,59^ To prepare these plates, *E. coli* strain OP50 was cultured in Luria-Bertani medium (25 g/L) at 37°C for overnight, and 75 μL of cell suspension was spread on each side of a NGM agar plate. Each test soil was arranged linearly in the central area of the soil-agar isolation plate, and offspring moving from the test soil to each side was counted. We expected that toxicity would be captured by fewer nematodes moving out of from the test soil into the fresh food resource. The data were expressed as a percentage (%) of average value of control group.

### Preparation of Extractable Additive Solution

Eactractable additive solutions were prepared from thirteen different microplastics (Table S1), and two methods were investigated using only liquid (method 1) and glass beads (method 2), respectively. In order to obtain extractable additive solution using method 1, 118.4 mg of each microplastic was placed into 10 mL-glass vials containing 3 mL of K-medium. Although microplastics either floated or sank depending on their different densities, hydrophobicity, or interaction with surface tension of microplastic, we did not attempt to immerse the particles in solution. The vials were maintained at 20 ± 2°C in the dark for 24 h, conditions similar to the soil toxicity test, and then the solutions were passed through a syringe filter (pore size 0.45 μm; D-76185, ROTILABO^®^, Carl Roth GmbH & Co., Karlsruhe, Germany). For method 2, glass beads (1-2 mm) were washed ten times using deionized water, and autoclaved at 121°C for 15 min, then dried at 60°C for 24 h. Each microplastic (118.4 mg) was added into 10 mL-glass vial containing 5 g of glass beads, and they were gently mixed using a spatula. Then 5 g of additional glass beads were placed on top of this bead-microplastic mixture, and then 3 mL of K-medium added. The microplastics were immersed in solution, similar to what the situation in soil water films inside of pores would be. The vials were maintained for 24 h at 20 ± 2°C in the dark, and the mixtures were moved to a 50 mL-syringe. The syringes were carefully pumped to obtain extractable solution, and the obtained solution was passed again through the glass bead-microplastic system in the syringe two times and then filtered using a syringe filter. As a result, we obtained 24 h-extractable additive solution from 0.04 microplastic mg/μl K-medium. We added 76 μL of this solution into each well of 24-well plate containing 0.3 g of soil, and the final concentration of our 24 h-extractable additive solution in soil (3.0 mg/0.3 g) corresponds to approximately 1% of microplastic in soil. The number of replicates was 4 (for method 1), 8 (HDPE, PET, PP, and PS fragments for method 2), and 4 (LDPE film and PAN fibers for method 2), respectively, and control sets were prepared with a matching equal number of replicates for every microplastic treatement set. Soil toxicity tests were performed, and negative control sets (without microplastics) were implemented for each method. The data were expressed as a percentage (%) of average value of each control group.

### Preparation of Microplastic with easily extractable materials removed

The additive-extracted microplastics were prepared using thirteen different microplastics (Table S1). We expected that microplastics can lose their harmful effects if the extractable additives are removed, and soil toxicity test was conducted using these extracted microplastics to test our hypothesis. We added 100 (for fragments and film) or 10 (for fibers) mg of each microplastic into 25 mL-glass vials containing 5 mL ethanol (96%), and these were maintained at 20 ± 2°C in the dark. We chose ethanol to remove extractable additives. Since the additives used in plastic products are mostly apolar, ethanol, which is slightly more apolar than water, could be better to extract from microplastic.^60^ We omitted stirring or shaking to avoid changing of size distributions of the microplastics. After 24 h, 4 mL of supernatant was removed, and 20 mL of deionized water was added to wash the microplastics. The suspensions were stabilized for 1 h, and 20 mL of supernatant (upper layer) or sub-natant (middle layer) was removed again by careful pipetting. This washing process was repeated three times, and then the vials containing microplastics were dried at 65°C for 24 h. In order to ensure that every available extractable additive is partitioned into the ethanol solution from the microplastic surface, these extraction procedures including ethanol-extraction and water-washing were repeated twice. These extracted microplastics with one and two extractions were mixed with soil, and the final concentration was determined as 1% (for fragments and film) or 0.1% (for fibers). We then added 0.3 g of each microplastic-laced soil into each well of a 24-well plate, and 76 μL of K-medium was poured into each well. The number of replicates was 8 (HDPE, PET, PP, and PS fragments for one time-extraction), 4 (LDPE film and PAN fibers for one time-extraction), and 4 (HDPE, PET, PP, and PS fragments for two times-extraction), respectively, and control sets were prepared with a matching equal number of replicates for every microplastic treatement set. Soil toxicity tests were performed, and negative controls (no microplastic added) were also implemented for the whole process. The data were expressed as a percentage (%) of average value of control group.

### Simulation of Wet-Dry Cycles in the Soil Environment

We selected LDPE film and PAN fibers as target materials for our extended experiment, because they are real plastic products circulating in commercial markets. In order to simulate wet-dry cycles in soil, 24-well plates containing each microplastic-laced soil were prepared using the same procedues as used for the soil toxicity test (0, 0.01, 0.1, and 1% for LDPE film; 0, 0.001, 0.01, and 0.1% for PAN fibers), and 76 μL of deionized water was added into each well (*n* = 4). We prepared three plates (first, second, and third) for both microplastics, and each plate was covered and maintained at 20 ± 2°C in the dark. After 6 days, all soil samples were dried, because water had evaporated. 76 μL of K-medium was added into each well of the first plate, and the same amount of deionized water was added into the second and third plates. The first plate was used for soil toxicity tests (6 days, first wet-dry cycle), and the others were maintained at 20 ± 2°C in the dark for an additional 6 days. Subsequently, the second plate was used for soil toxicity tests (12 days, second wet-dry cycle), and the third plate after an additional 6 days (18 days, third wet-dry cycle). Negative controls (no microplastic added) were also prepared, and handled the same way.

### Statistical Analyses

Data were analyzed using the SPSS statistical software (Ver. 24.0, SPSS Inc., Chicago, IL, USA). One-way analysis for variance (ANOVA) and Turkey’s tests were conducted to determine the significance (*p* < 0.05) of multiple comparisons.

## RESULTS

### Effects of Microplastics on Nematodes in Soil

*C. elegans* showed vigorous reproductive activity in our soils, and an average value of offspring number per replicate was calculated as 171 ± 50 worms (*n* = 26) in control soil, which is comparable to international standards.^61^ Microplastic exposure showed that HDPE and PS fragments induce a significant effect on nematodes at the higher concentration, 1% (Figure 1A and 1D). By comparison, PET fragments started to be significantly harmful at 0.1% (Figure 1B), and PP influenced nematode offspring just for microplastics smaller than 250 μm, at the higher concentration of 1% (Figure 1C). There was no effect of the LDPE film (Figure 1E), and PAN fibers induced significant reproduction decrease at 0.1% (Figure 1F). In summary, microplastic mostly influenced nematodes at 1% concentration, and the number of offspring decreased to 78-80% (for PP and PAN) and 56-68% (HDPE, PET, and PS) compared with the control group. The toxic effects appear to increase in the order of LDPE film < PP fragments < HDPE fragments ≈ PS fragments < PET fragments ≈ PAN fibers. PP fragments were the only plastic inducing a size-dependent effect.

**Figure 1.**
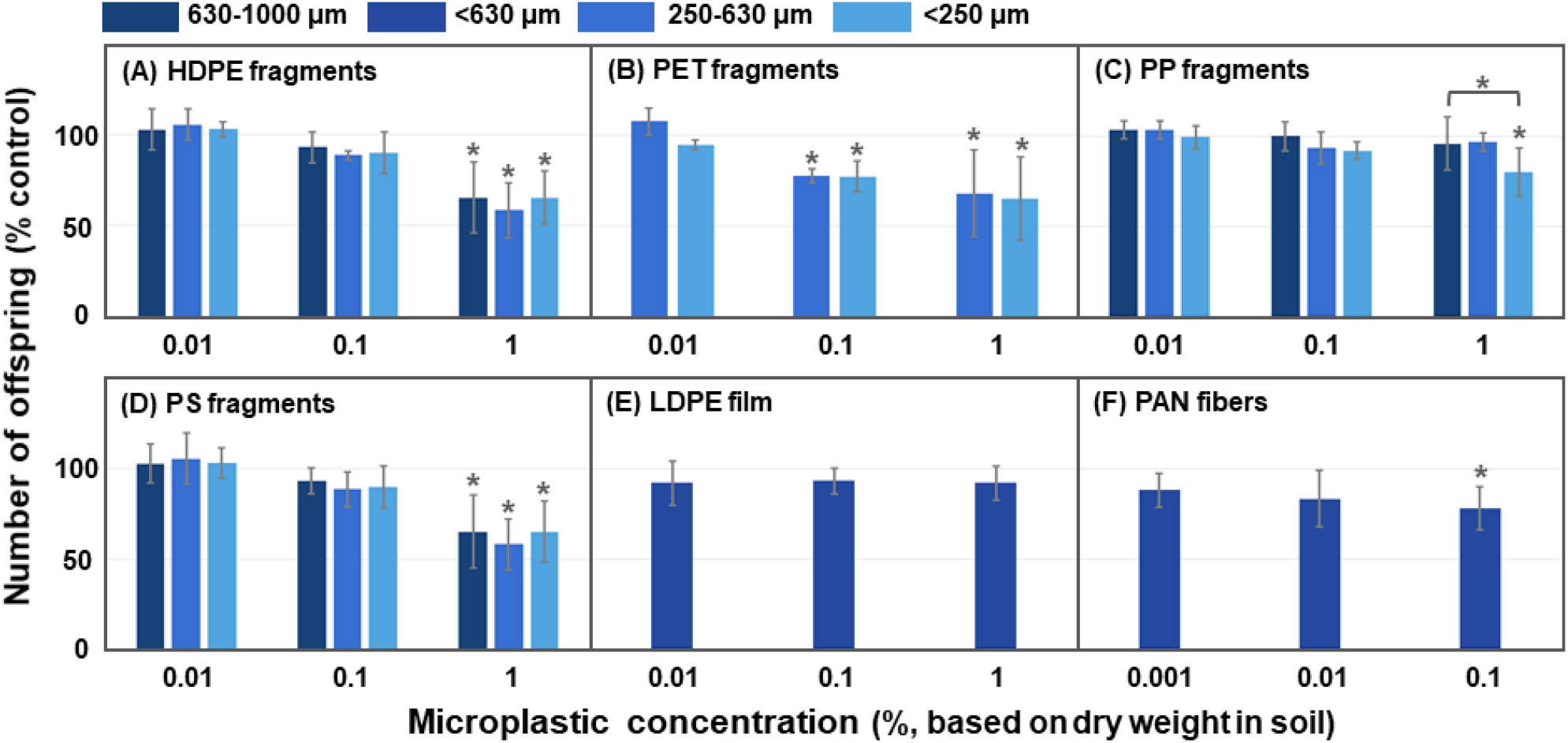
The offspring number of *C. elegans* exposed to (A) high-density polyethylene (HDPE) fragments, (B) polyethylene terephthalate (PET) fragments, (C) polypropylene (PP) fragments, (D) polystyrene (PS) fragments, (E) low-density polyethylene (LDPE) film, and (F) polyacrylicnitrile (PAN) fibers in soil. Each microplastic contains one to three different size ranges (<250, 250-630, <630, and 630-1000 μm), and test concentrations are expressed as percentage (%) based on dry weight in soil. All data are normalized to each control group, and error bars indicate standard deviations. The asterisks (*) indicate significant (*p* < 0.05) differences compared to the control or the other different sizes.

### Effects of Extractable Additive Solutions

The 24 h-extractable additive solution was acquired using two methods (method 1 with liquid and 2 with glass beads). Average values of offspring were 174 ± 24 (*n* =8) and 161 ± 12 (*n* = 12) worms in each negative control soil (no microplastic added) for methods 1 and 2, respectively. As shown in Figure 2A, additives extracted using method 1 had no effects, while method 2 led to a significant percentage decline of the number of offspring to 79 ± 11 (HDPE fragments, 630-1000 μm), 84 ± 5 (HDPE fragments, 250-630 μm), 84 ± 7 (HDPE fragments, <250 μm), 80 ± 5 (PET fragments, 250-630 μm), 84 ± 6 (PET fragments, <250 μm), 84 ± 10 (PP fragments, <250 μm), 77 ± 12 (PS fragments, 630-100 μm), 83 ± 8 (PS fragments, 250-630 μm), and 75 ± 9 (PS fragments, <250 μm), compared with the control group (Figure 2B). There were no significant effects of larger PP fragments (630-1000 and 250-630 μm), LDPE film, and PAN fibers. These toxicity trends were very similar to the results presented in Figure 1.

**Figure 2.**
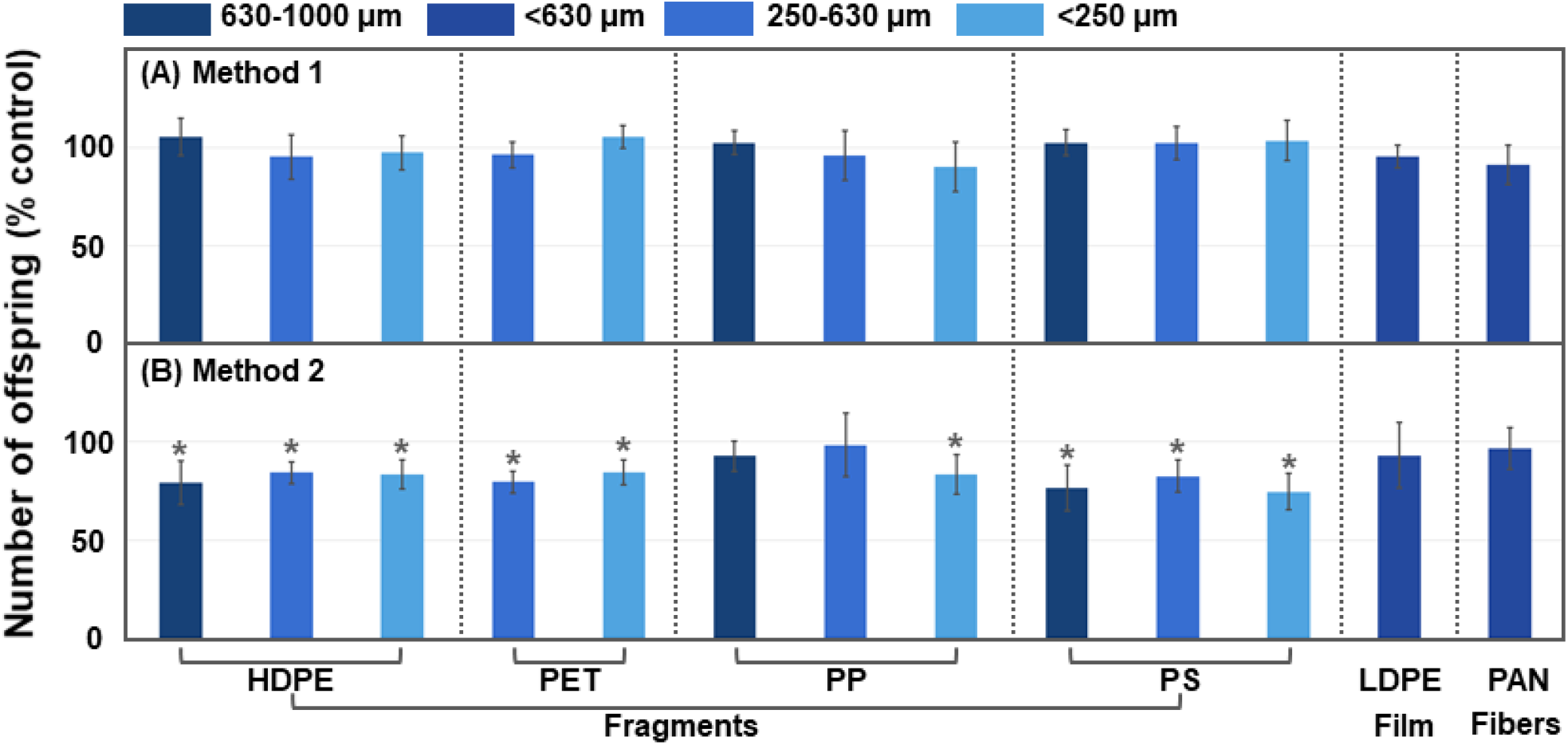
The offspring number of *C. elegans* exposed to 24 h-extractable additive solutions which are obtained by (A) method 1 (only liquid) and (B) method 2 (glass beads). Each 24 h-extractable additive solution was prepared using high-density polyethylene (HDPE) fragments, polyethylene terephthalate (PET) fragments, polypropylene (PP) fragments, polystyrene (PS) fragments, low-density polyethylene (LDPE) film, and polyacrylicnitrile (PAN) fibers, and final concentrations were determined with an approximate level of additive concentration from 1% or 0.1% (PAN fibers) based on dry weight in soil (see methods). Each microplastic contains one to three different size ranges (<250, 250-630, <630, and 630-1000 μm). All data are normalized to each control group, and error bars indicate standard deviations. The asterisks (*) indicate significant (*p* < 0.05) differences compared with the control.

### Effects of the Additive-Extracted Microplastics

Average values of offspring were 166 ± 35 (*n* =12) and 165 ± 22 (*n* = 4) in each negative control experiencing extraction procedures (without microplastics) for one and two times, respectively. Extracting the microplastics once (Figure 3A) led to the disappearance of the toxic effects of PP and PS fragments. Still, HDPE fragments (250-630 and <250 μm) significantly reduced the offspring number to 80 ± 10 and 80 ± 8% compared to the control, respectively. PET fragments (250-630 and <250 μm) also still showed a toxic effect to 86 ± 5 and 80 ± 6% of the control, respectively. When the extraction procedures were repeated twice, there were no more toxic effects for any the microplastics (Figure 3B).

**Figure 3.**
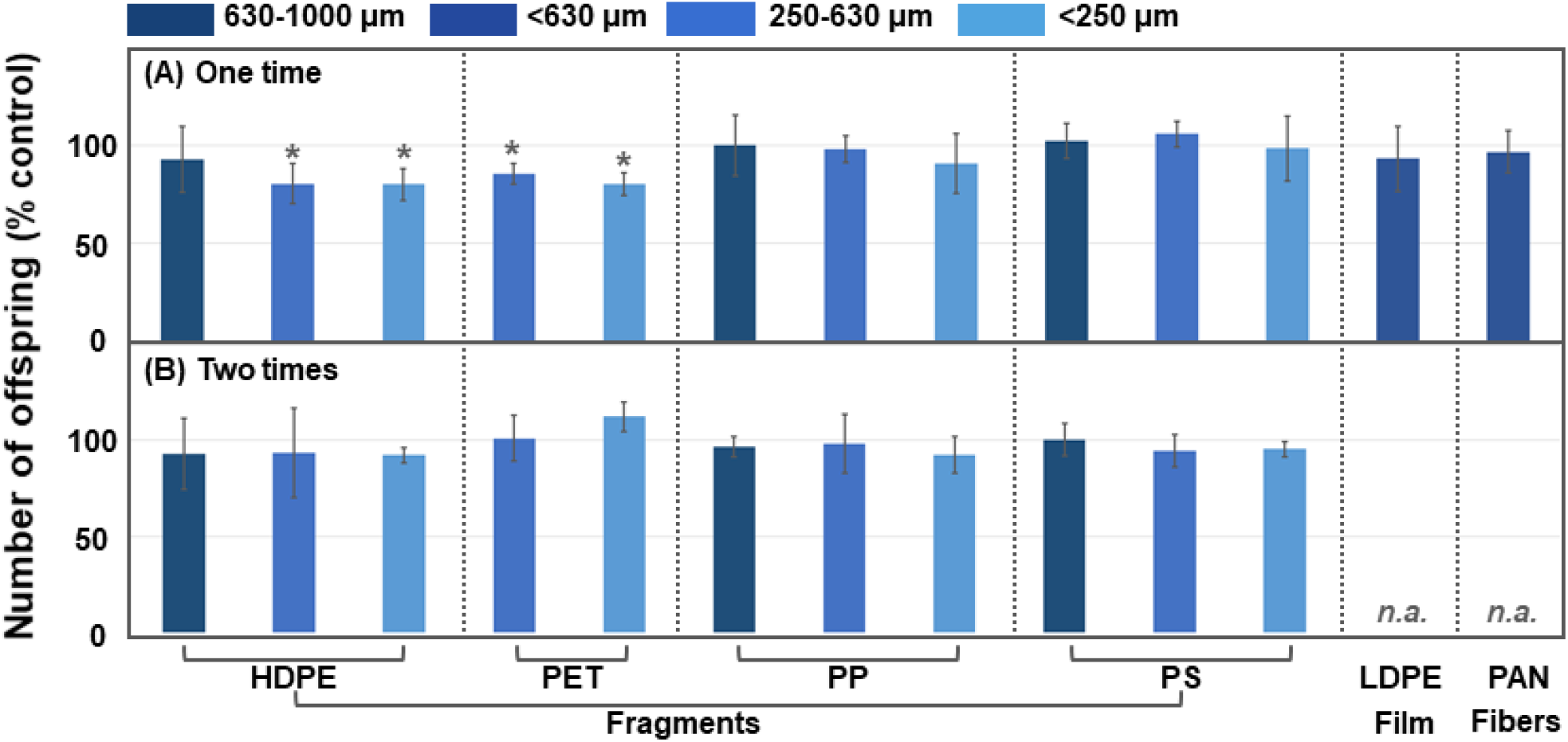
The offspring number of *C. elegans* exposed to extracted microplastic for (A) one extraction round, and (B) two rounds of extraction. Each extracted microplastic was prepared using high-density polyethylene (HDPE) fragments, polyethylene terephthalate (PET) fragments, polypropylene (PP) fragments, polystyrene (PS) fragments, low-density polyethylene (LDPE) film, and polyacrylicnitrile (PAN) fibers, and final concentrations were determined as 1% or 0.1% (PA fibers) based on dry weight in soil. Each microplastic contains one to three different size ranges (<250, 250-630, <630, and 630-1000 μm). All data are normalized to each control group, and error bars indicate standard deviations. The asterisks (*) indicate significant (*p* < 0.05) differences compared with the control.

### Simulation of Wet-Dry Cycle in Soil Environment

After one wet-dry cycle (6 days), the number of offspring significantly decreased for the LDPE film at all concentrations (0.01 to 1%), and average values were 70 ± 11, 69 ± 17, and 41 ± 8% compared to the control, respectively (Figure 4A). Toxic effects were intensified to 43 ± 9, 41 ± 5, 34 ± 8% at each concentration after two wet-dry cycles (12 days) (Figure 4B), and these effects were maintained at 42-58% after three wet-dry cycles (18 days) (Figure 4C). When LDPE film was extracted before the experiment, significant effects did not appear until two wet-dry cycles (Figures 4A and B), and 39% of the reproduction level was observed after three wet-dry cycles (Figure 4C). In the case of PAN fibers, the number of offspring significantly decreased at 0.01 and 0.1% after one wet-dry cycle (6 days) with average values of 74 ± 12 and 46 ± 11 compared to the control, respectively (Figure 4D). These effects were intensified at all concentrations (0.001 to 0.1%) after two and three wet-dry cycles, and 42 to 57% of the reproduction level was found (Figure 4E and F). When the PAN fibers were extracted, significant effects started to appear after two wet-dry cycles (Figure 4E), and 49-53% of the reproduction level was maintained until three wet-dry cycles (Figure 4F). Figure 5 shows that the toxic effects of LDPE film (1%) and PAN fibers (0.1%) increased as a function of the repetition of the wet-dry cycle towards a plateau of around 34-56%. Extraction procedures slowed down the appearance of toxic effects, but both treated and non-treated microplastics showed a trend to the same plateau after three wet-dry cycles, at 18 days.

**Figure 4.**
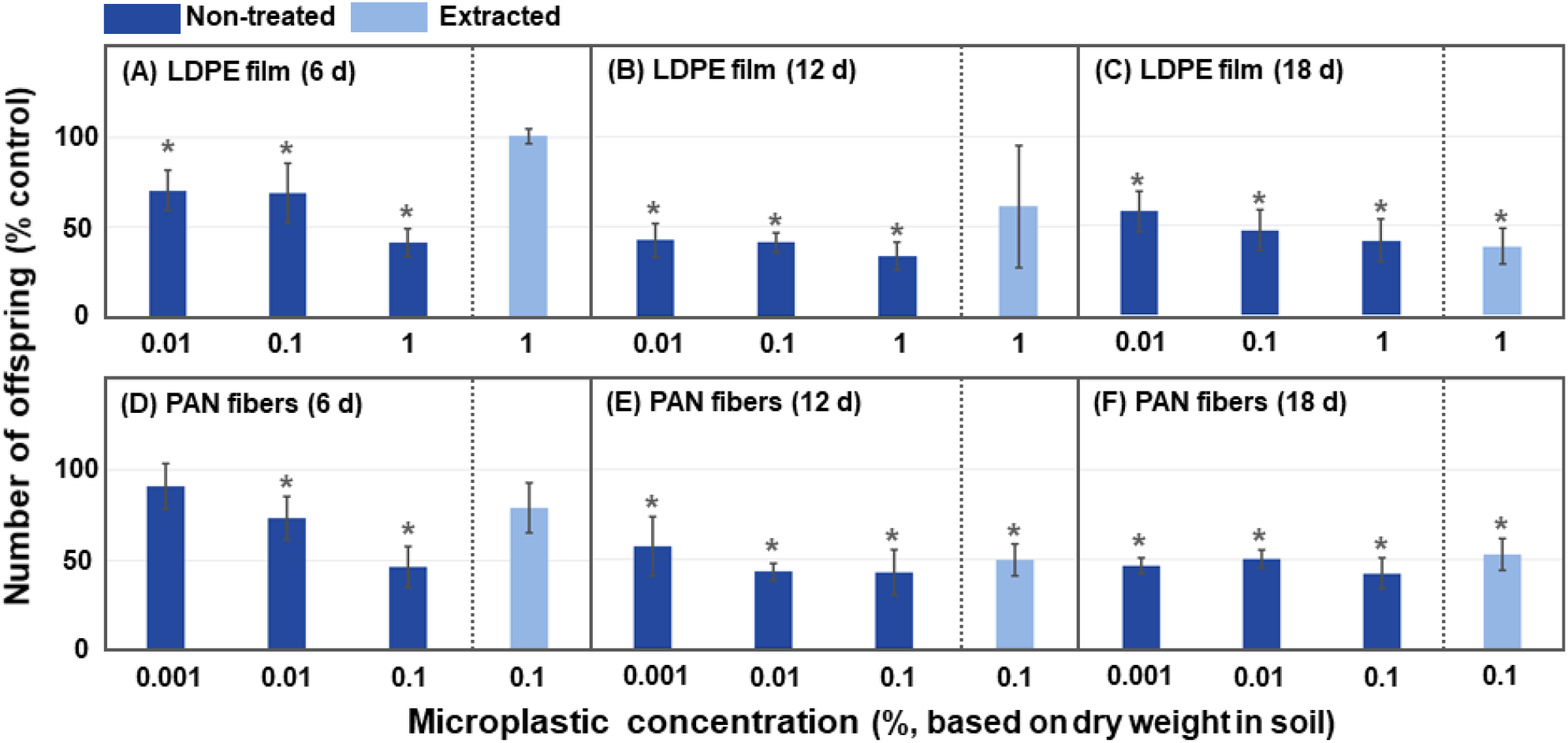
The offspring number of *C. elegans* exposed to (A) LDPE film and (B) PAN fibers in soil. Each soil was maintained for (A,B) 6 days, (B,E) 12 days, and (C,F) 18 days, and experienced one wet-dry cycle every 6 days. Test concentrations are expressed to percentage (%) based on dry weight in soil. All data are normalized to each control group, and error bars indicate standard deviations. The asterisks (*) indicate significant (*p* < 0.05) differences compared with the control.

**Figure 5.**
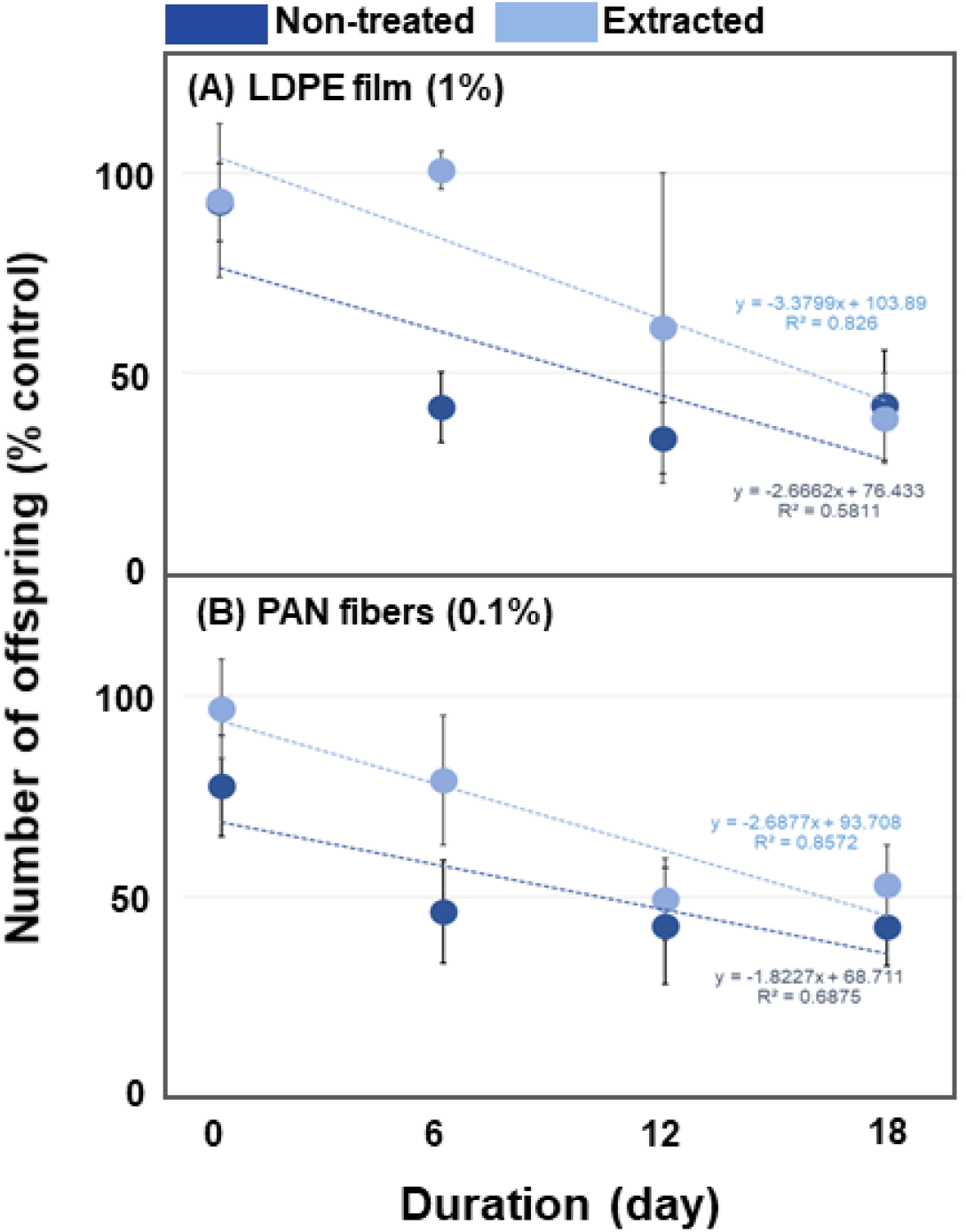
The offspring number of *C. elegans* exposed to (A) LDPE film (1%) and (B) PAN fibers (0.1%) in soil. Figure depicts the same data as Figure 4, but here focus on time-dependent changes with linear trend lines. Each soil was maintained for 6 to 18 days, and experienced one wet-dry cycle every 6 day. Test concentrations are expressed to percentage (%) based on dry weight in soil. All data are normalized to each control group, and error bars indicate standard deviations.

## DISCUSSION

### Effects of Microplastics on Nematodes in Soil

*C. elegans* is one of the most extensively studied species for microplastic toxicity research, and 26 scientific papers have been published until March 31, 2020 (Table S2). These studies provide reliable, initial information aiding our understanding of microplastic toxicity on nematodes, but they also left open many important points. Notably, most of these studies except only four papers^29,62–64^
 have adopted spherical PS particles as target material, and only two papers are utilizing field-collected or secondary-treated particles instead of purchased beads or pellets.^62,64^ Although *C. elegans* has been suggested as a standard soil test species,^61,65^ there is only one study conducting tests in soil media,^28^ and 25 studies were performed using liquid media such as K-medium and M9 buffer solution (Table S2). On the other hand, six papers report size-dependent inhibitory effects of microplastic on *C. elegans*,^27–30,66,67^ showing a tendency toward toxic effects that can be increased by smaller sizes in ranges of 0.05 to μm,^27^ 0.1 to 6.0 μm,^67^ and 0.1 to 5.0 μm.^29^ Lei et al.^30^ reported that effects of microplastics in this smaller size range might not be linear, since the intermediate-sized group (1.0 μm) had the lowest survival rate, compared to smaller and larger sizes (0.1 to 0.5 and 2.0 to 5.0 μm), and Muller et al.^67^ found toxicity to increase in >10 μm-size range. In our study, we used a larger size range (around 250 to 1000 μm) than previous work (0.05 to 6 μm). Since the edible size of microplastic by nematode species is ≤3.4 μm,^66,67^ we avoided that the nematodes fed on microplastics and followed just the influence of the potential leachates in the number of nematodes offspring. PP microplastics had a size-dependent effect, with toxicity only apparent for its smaller size range (<250 μm). Regarding the concentration, our results showed that most of the microplastics had toxic effects after 24 h when present in higher concentrations in the soil (Figure 1). HDPE, PET, and PS presented concentration-effect toxicity since only the higher concentrations presented toxic effects on nematode offspring numbers.

### Production of Extractable Additive Solutions and their effects

Toxicity of microplastics is often associated with the pollutants they sorb during exposure to the environment and the chemicals used as additives leached during useful life and after being discarded.^68^ Regarding the additives, they are moving through the bulk of the microplastic particle until they eventually reach the surface, where they might stay or migrate to the surrounding medium.^69^ Our work is focused on chemicals leaching and on the concept of K_D_ to better understand the ecotoxicity of microplastics. The K_D_ is the equilibrium of a chemical between two immiscible media; in this study, it is between the surface of the microplastic and the aqueous environment within the soil.^70^ When the K_D_ is high, this means that additives will interact more with the apolar part, even though a small portion will migrate into the aqueous medium (Figure 6A). When the K_D_ is low, this means that most of the chemicals will be released into the aqueous surrounding matrix, even if a small portion still adheres to the surface of the microplastics (Figure 6B). Finally, the real picture for plastics typically means the presence of a mix of additives,^71^ with a range of K_D_ (Figure 6C). In such a mixture, it is likely to have the major fraction of the chemicals with higher K_D_ mostly on the surface of the microplastics and the major part of the chemicals with lower K_D_ in the aqueous environment.^72^ Since the microplastics used in this study had no history of exposure to the environment - thus no sorption of pollutants - and the sizes used were not small enough to be ingested by nematodes, the most likely explanation of toxic effects, expressed as a reduction in nematode offspring, is leached chemicals from the microplastics to the soil. To evaluate this hypothesis, we used an extract produced under very mild conditions for the migration of an apolar additive: 24 hours of contact with water for the leaching and then using this solution for the toxicity test.

**Figure 6.**
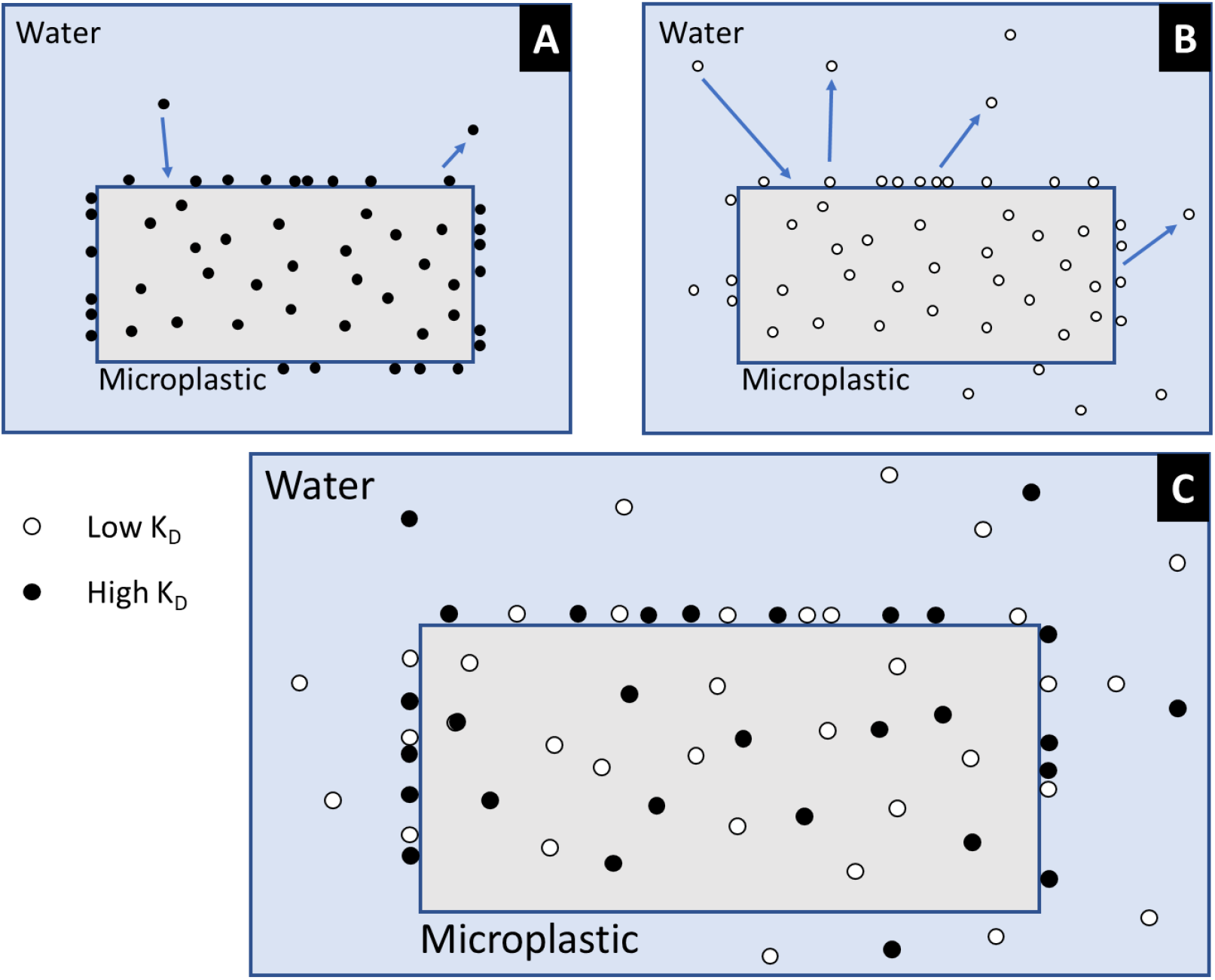
Scheme for the relative abundance of chemicals with (A) a high partition constant (K_D_), (B) low K_D_, and (C) a mixture of chemicals with a range of K_D_. The arrows in A and B represent the dynamic behavior chemical movement.

The outcomes of leaching tests depend heavily on methodology,^44^ and several experiments have been conducted to simulate various leaching environments under laboratory conditions such as shaking,^41,42^ static maintenance,^40,44^ and standard leaching method.^43^ These approaches are based on the concept of leaving microplastics afloat because this is likely close to natural exposure conditions in an aquatic environment.^44^ However, this exposure scenario is not fully applicable to the soil environment, and a direct application of standard leaching methods including soil column test,^73,74^ batch test using the liquid to solid ratio,^75,76^ and up-flow percolation test^77^ are difficult due to a wide variation of plastic characteristics. Also, the standard leaching tests have focused on traditional pollutants such as metals and organic chemicals, and these materials have been well characterized in terms of basic information on which factors control leachability.^78^ Since we have no such knowledge about microplastic in soil, we should be cautious about determining experimental procedures. In this study, we assessed two different methods for the chemicals leaching to the water: 1) floating in a liquid to emulate the conditions in aquatic bodies in nature; and 2) using glass beads to keep microplastics immersed in water. While there was no effect using the 24 h-extractable solution obtained by floating microplastics in water (Figure 2A), the number of nematode offspring significantly decreases when using the 24 h-extractable solution prepared using glass beads (Figure 2B). The more efficient migration was likely due to the better interaction with water since the microplastics were in complete contact with the water while the floating microplastics were only partially in contact with water, with a lower interface area.

### Effects of Additive-Extracted Microplastic

After determining the protocol for a more efficient migration of the additives, using glass beads, we tested if the additives were indeed the toxicity source, trying to remove them from the microplastic particles. Since the additives were successfully removed even with water, we tested the extraction with ethanol, a polar solvent, but less polar than water. The higher the ethanol content, the more effective is the migration of organic chemicals from plastics to the solution. Although a K_D_ value depends on properties of target chemical migrants and plastics,^79–82^ we believe that as a general rule, ethanol can promote the migration of apolar additives because it is less polar than water. For example, K_LDPE/95%-ethanol_ at 60°C is 775-times lower than in 50%-ethanol at the same temperature,^69^ and K_95%-ethanol/PET_ at 20°C is 3 to 4-times higher than in water.^82^ After one extraction, we observed that the significant effects of HDPE and PET fragments remained (Figure 3A), but all the other microplastics no longer had toxic effects. To confirm the effect, we extracted once more (Figure 3B), and the result was no toxic effect of any microplastics tested irrespective of concentrations and shapes. We concluded, therefore, that the toxic effects of microplastic are mainly caused by the 24 h-extractable additives from the microplastics.

### Simulation of Wet-Dry Cycles in the Soil Environment

The diffusion of chemicals through the bulk of the plastic proceeds until they reach the surface and migrate to the other medium in a proportion regulated by the K_D_. The kinetics of the diffusion influences the amount of chemicals on the surface, and thus the migration to the environment. To determine the duration of the whole process of diffusion and migration, we tested the time needed to produce toxic effects on the nematodes. We used the samples purchased from the market, LDPE film, and PAN fibers. Desorption of hydrophobic organic pollutants from plastics is generally slow, and the leaching rate of chemical additives from plastic into water depends on time.^83–85^ For example, the desorption half-lives of polychlorinated biphenyls from PE pellets are estimated to be 14 days to 210 years,^83^ and the leaching rate of brominated diphenyl ethers-209 from HDPE plate is calculated as 2.1 × 10^5^ ng/m^2^/d.^86^ Chemicals keep leaching from microplastics until depletion.^85^ When the microplastics are present in a soil system with low diffusivity or in a closed system (like a laboratory experiment), we can expect an increasing concentration of the chemicals leached. In this study, we expected that the toxic effects of LDPE film and PAN fibers can be intensified by repeating a wet-dry cycle in soil. Since the test duration for extractable additives was only 24 h, according to soil toxicity test conditions, there is a high possibility of toxicity increase if microplastics are maintained in simulated soil conditions for more extended periods. Our expectation was correct, and we found that these effects plateaued with a similar decreasing level until 18 days (Figure 4). Extracted microplastics showed a relatively slower increasing trend of toxic effects compared to non-treated ones (Figure 5). Our result indicates that the extractable additives from plastics can be more harmful when they are maintained in soil environments for a longer period than those used in typical testing protocols,^41–44^ and toxic effects can occur at a relatively low concentration like 0.01% (100 mg kg^−1^) for LDPE film and 0.001% (10 mg/kg) for PAN fibers. Since 8 to 67500 mg kg^−1^ of microplastic can be detected in the soil environment,^9–11^ nematode populations would be expected to be affected given the microplastic concentrations we tested here.

We conducted a simple ecotoxicological protocol using the concept of diffusion and migration of chemical additives from microplastics. Although our study was performed on a small scale taking a more phenomenological approach, our ecotoxicological tests provide clear evidence that microplastic toxicity in soil is linked with their characteristics and extractable additives. This study is the first to estimate microplastic levels inducing toxic effects on nematodes in the soil system, uncovering particle characteristics and the crucial role of extractrable additives. Our results strongly suggest that future test consider microplastic additives as a key explanatory variable.

## ASSOCIATED CONTENT

### Supporting Information

The Supporting information is available free.

■ Methods: FTIR spectra of each target MP (Figure S1), List of target plastic materials (Table S1)
■ Result and Discussion: List of previous studies reporting microplastic toxicity on the nematode *C. elegans* (Table S2)

## ACKNOWLEDGEMENTS

This work was supported by a post-doctoral grant from the National Research Foundation of Korea funded by the Ministry of Science, ICT, and Future Planning (2019R1A6A3A03031386). MCR acknowledges support from an ERC Advanced Grant (grant no. 694368). WRW acknowledges a Capes-Humboldt Research Fellowship (1203128-BRA-HFSTCAPES-E-Finance code 001). We also thank to Dr. Moises Sosa Hernandez for providing test soil.

## REFERENCES

1. Andrady, A. L. Microplastics in the marine environment. Mar. Pollut. Bull. 2011, 62, 1596–1605.

2. Lambert, S.; Wagner, M. Microplastics are contaminants of emerging concern in freshwater environments: An Overview. In Freshwater Microplastics. Emerging Environmental Contaminants?; Wagner, M., Lambert, S., Eds.; Springer, Gewerbestrasse, Switzerland, 2018; pp. 1–23.

3. Koelmans, A. A.; Nor, N. H. M.; Hermsen, E.; Kooi, M.; Mintenig, S. M.; de France, J. Microplastics in freshwaters and drinking water: Critical review and assessment of data quality. Water Res. 2019, 155, 410–422.

4. Wang, J.; Liu, X.; Li, Y.; Powell, T.; Wang, X.; Wang, G.; Zhang, P. Microplastics as contaminants in the soil environment: A mini-review. Sci. Total Environ. 2019, 691, 848–857.

5. Chen, G.; Feng, Q.; Wang, J. Mini-review of microplastics in the atmosphere and their risks to humans. Sci. Total Environ. 2020. 703, 135504.

6. Pivokonsky, M.; Cermakova, L.; Novotna, K.; Peer, P.; Cajthaml, T.; Janda, V. Occurrence of microplastics in raw and treated drinking water. Sci. Total Environ. 2018, 643, 1644–1651.

7. Horton, A. A.; Walton, A.; Spurgeon, D. J.; Lahive, E.; Svendsen, C. Microplastics in freshwater and terrestrial environments: evaluating the current understanding to identify the knowledge gaps and future research priorities. Sci. Total Environ. 2017, 586, 127–141.

8. Bläsing, M.; Amelung, W. Microplastics in soil: Analytical methods and possible sources. Sci. Total Environ. 2018, 612, 422–435.

9. Fuller, S.; Gautam, A. A procedure for measuring microplastics using pressurized fluid extraction. Environ. Sci. Technol. 2016, 50, 5774–5780.

10. Scheurer, M.; Bigalke, M. Microplastics in Swiss floodplain soils. Environ. Sci. Technol. 2018, 52, 3591–3598.

11. Liu, M.; Lu, S.; Song, Y.; Lei, L.; Hu, J.; Lv, W.; Zhou, W.; Cao, C.; Shi, H.; Yang, X.; He, D. Microplastic and mesoplasitc pollution in farmland soils in suburbs of Shanghai, China. Environ. Pollut. 2018, 242, 855–862.

12. Backhaus, T.; Wagner, M. Microplastic in the environment: Much ado about nothing? A debate. Global Challenges. 2018, 190022.

13. Li, J.; Liu, H.; Chen, P. Microplastics in freshwater systems: A review on occurrence, environmental effects, and methods for microplastics detection. Water Res.. 2018, 137, 362–374.

14. de Souza Machado, A. A.; Lau, C. W.; Kloas, W.; Bergmann, J.; Bachelier, J. B.; Faltin, E.; Becker, R.; Görlich, A. S.; Rillig, M. C. Microplastics can change soil properties and affect plant performance. Environ. Sci. Technol. 2019, 53, 6044–6052.

15. Huerta Lwanga, E.; Gertsen, H.; Gooren, H.; Peters, P.; Salanki, T.; van der Ploeg, M.; Besseling, E.; Koelmans, A. A.; Geissen, V. Microplastics in the terrestrial ecosystem: implications for *Lumbricus terrestris* (Oligochaeta, Lumbricidae). Environ. Sci. Technol. 2016, 50, 2685–2691.

16. Ju, H.; Zhu, D.; Qiao, M. Effects of polyethylene microplastics on the gut microbial community, reproduction and avoidance behaviors of the soil springtail, *Folsomia candida*. Environ. Pollut. 2019, 247, 890–597.

17. Qi, Y.; Yang, X.; Pelaez, A. M.; Huerta Lwanga, E.; Beriot, N.; Gertsen, H.; Garbeva, P.; Geissen, V. Macro- and micro- plastics in soil-plant system: Effects of plastic mulch film residues on wheat (*Triticum aestivum*) growth. Sci. Total Environ. 2018, 645, 1048–1056.

18. Awet, T. T.; Kohl, Y.; Meier, F.; Straskraba, S.; Grün, A.-L.; Ruf, T.; Jost, C.; Drexel, R.; Tunc, E.; Emmerling, C. Effects of polystyrene nanoparticles on the microbiota and functional diversity of enzymes in soil. Environ. Sci. Eur. 2018, 30, 11.

19. Huang, Y.; Zhao, Y.; Wang, J.; Zhang, M.; Jia, W.; Qin, X. LDPE microplastic films alter microbial community composition and enzymatic activities in soil Environ. Pollut. 2019, 254, 112983.

20. Kooi, M.; Kolemans, A. A. Simplifying microplastic via continuous probability distributions for size, shape, and density. Environ. Sci. Technol. 2019, 6, 551–557.

21. Rillig, M. C.; Lehmann, A.; Ryo, M.; Bergmann, J. Shaping up: towards considering the shape and form of pollutants. Environ. Sci. Technol. 2019, 53, 7925–7926

22. Choi, J. S.; Jung, Y.-J.; Hong, N.-H.; Hong, S. H.; Park, J.-W. Toxicological effects of irregularly shaped and spherical microplastics in a marine teleost, the sheepshead minnow (*Cyprinodon variegatus*). Mar. Pollut. Bullet. 2018, 129, 231–240.

23. Gray, A. D.; Weinstein, J. E. Size- and shape-dependent effects of microplastic particles on adult daggerblade grass shrimp (*Palaemonetes pugio*). Environ. Toxicol. Chem. 2017, 36, 301–3080.

24. Jeong, C. B.; Won, E.-J.; Kang, H.-M.; Lee, M.-C.; Hwang, D.-S.; Hwang, U.-K.; Zhou, B.; Souissi, S.; Lee, S.-J.; Lee, J.-S. Microplastic size-dependent toxicity, oxidative stress induction, and p-JNK and p-p38 activation in the monogonont rotifer (*Brachionus koreanus*). Environ. Sci. Technol. 2016, 50, 8849–8857.

25. Lagarde, F.; Olivier, O.; Zanella, M.; Daniel, P.; Hiard, S.; Caruso, A. Microplastic interactions with freshwater microalgae: Hetero-aggregation and changes in plastic density appear strongly dependent on polymer type. Environ. Pollut. 2016, 215, 331–339.

26. Jiang, X.; Chen, H.; Liao, Y.; Ye, Z.; Li, M.; Klobučar, G. Ecotoxicity and genotoxicity of polystyrene microplastics on higher plant *Vicia faba*. Environ. Pollut. 2019, 250, 831–838.

27. Kim, H. M.; Lee, D.-K.; Long, N. P.; Kwon, S. W.; Park, J. H. Uptake of nanopolystyrene particles induces distinct metabolic profiles and toxic effects in *Caenorhabditis elegans*. Environ. Pollut. 2019, 246, 578–586.

28. Kim, S. W.; Kim, D.; Jeong, S.-W.; An, Y.-J. Size-dependent effects of polystyrene plastic particles on the nematode *Caenorhabditis elegans* as related to soil physicochemical properties. Environ. Pollut. 2020, 258, 113740.

29. Lei, L.; Wu, S.; Lu, S.; Liu, M.; Song, Y.; Fu, Z.; Shi, H.; Raley-Susman, K. M.; He, D. Microplastic particles cause intestinal damage and other adverse effects in zebrafish *Danio rerio* and nematode *Caenorhabditis elegans*. Sci. Total Environ. 2018, 619–620, 1–8.

30. Lei, L.; Liu, M.; Song, Y.; Lu, S.; Hu, J.; Cao, C.; Xie, B.; Shi, H.; He, D. Polystyrene (nano)microplastics cause size-dependent neurotoxicity, oxidative damage and other adverse effects in *Caenorhabditis elegans*. Environ. Sci. Nano 2018, 5, 2009.

31. Wan, Y.; Wu, C.; Xue, Q.; Hui, X. Effects of plastic contamination on water evaporation and desiccation cracking in soil. Sci. Total Environ. 2019, 654, 576–582.

32. Koelmans, A. A.; Bakir, A.; Burton, A.; Janssen, C. R. Microplasatic as a vector for chemicals in the aquatic environment: Critical review and model-supported reinterpretation of empirical studies. Environ. Sci. Technol. 2016, 50, 3315–3326.

33. Kwon, J.-H.; Chang, S.; Hong, S. H.; Shim, W. J. Microplastic as a vector of hydrophobic contaminants: Importance of hydrophobic additives. Integ. Environ. Asses. 2017, 13, 494–499.

34. Hahladakis, J. N.; Velis, C. A.; Weber, R.; Iacovidou, E.; Purnell, P. An overview of chemical additives present in plastics: Migration, release, fate and environmental impact during their use, disposal and recycling. J. Hazard. Mater. 2018, 344, 179–199.

35. Ügdüler, S.; van Geem, K. M.; Roosen, M.; Delbeke, E. I. P.; de Meester, S. Challenges and opportunities of solvent-based additive extraction methods for plastic recycling. Waste Manag. 2020, 104, 148–182.

36. Wei, W.; Huang, Q.-S.; Sun, J.; Wang, J.-Y.; Wu, S.-L., Ni, B.-J. Polyvinyl Chloride Microplastics Affect Methane Production from the Anaerobic Digestion of Waste Activated sludge through leaching toxic bisphenol-A. Environ. Sci. Technol. 2019, 53, 2509–5217.

37. Fikarová, K.; Cocovi-Solberg, D. J.; Rosende, M.; Horstkotte, B.; Sklenárová, H.; Miró, M. A flow-based platform hyphenated to on-line liquid chromatography for automatic leaching tests of chemical additives from microplastics into seawater. J. Chromatogr. A 2019, 1602, 160–167.

38. Oehlmann, J.; Schulte-Oehlmann, U.; Kloas, W.; Jagnytsch, O.; Lutz, I.; Kusk, K. O.; Wollenberger, L.; Santos, E. M.; Paull, G. C.; van Look, K. J. W.; Tyler, C. R. A critical analysis of the biological impacts of plasticizers on wildlife. Phil. Trans. R. Soc. B 2009, 364, 2047–2062.

39. Lithner, D.; Nordensvan, I.; Dave, G. Comparative acute toxicity of leachates from plastic products made of polypropylene, polyethylene, PVC, acrylonitrile–butadiene–styrene, and epoxy to *Daphnia magna*. Environ. Sci. Pollut. Res. 2012, 19, 1763–1772.

40. Schrank, I.; Trotter, B.; Dummert, J.; Scholz-Bottcher, B. M.; Loder, M. G. J.; Laforsch, C. Effects of microplastic particles and leaching additive on the life history and morphology of *Daphnia magna*. Environ. Pollut. 2019, 255, 113233.

41. Chae, Y.; Hong, S. H.; An, Y.-J. Photosynthesis enhancement in four marine microalgal species exposed to expanded polystyrene leachate. Ecotoxicol. Environ. Safe. 2020, 189, 109936.

42. Luo, H.; Xiang, Y.; He, D.; Li, Y.; Zhao, Y.; Wang, S.; Pan, X. Leaching behavior of fluorescent additives from microplastics and the toxicity of leachate to *Chlorella vulgaris*. Sci. Total Environ. 2019, 678, 1–9.

43. Bejgran, S.; MacLeon, M.; Bogdal, C.; Breitholtz, M. Toxicity of leachate from weathering plastics: An exploratory screening study with *Nitocra spinipes*. Chemosphere 2015, 132, 114–119.

44. Gandara e Silva, P. P.; Nobre, C. R.; Resaffe, P.; Pereira, C. D. S.; Gusmao, F. Leachate from microplastics impairs larval development in brown mussels. Water Res.. 2016, 106, 364–370.

45. Lee, H.; Shim, W. J.; Kwon, J.-H. Sorption capacity of plastic debris for hydrophobic organic chemicals. Sci. Total. Environ. 2014, 470–471, 1545–1552.

46. Smedes, F.; Geertsma, R. W.; van Der Zande, T., Booij, K. Polymer-water partition coefficients of hydrophobic compounds for passive sampling: Application of cosolvent models for validation. Environ. Sci. Technol. 2009, 43, 7047–7054.

47. Besseling, E.; Quik, J. T. K.; Sun, M.; Koelmans, A. A. Fate of nano- and microplastic in freshwater systems: A modeling study. Environ. Pollut. 2017, 220, 540–548.

48. Porter, A.; Lyons, B. P.; Galloway, T. S.; Lewis, C. Role of marine snows in microplastic fate and bioavailability. Environ. Sci. Technol. 2018, 52, 7111–7119.

49. Maaß, S.; Daphi, D.; Lehmann, A.; Rillig, M. C. Transport of microplastics by two collembolan species. Environ. Pollut. 2017, 225, 456–459.

50. O’Connor, D.; Pan, S.; Shen, Z.; Song, Y.; Jin, Y.; Wu, W.-M.; Hou, D. Microplastics undergo accelerated vertical migration in sand soil due to small size and wet-dry cycles. Environ. Pollut. 2019, 249, 527–534.

51. Kim, H. S.; Kim, S.-K.; Kim, J.-G.; Lee, D. S. Are the ratios of the two concentrations at steady state in the medium pairs of air-water, air-soil, water-soil, water-sediment, and soil-sediment? Sci. Total Environ. 2016, 553, 52–59.

52. Kim, H. S.; Lee, D. S. Proximity to chemical equilibria among air, water, soil, and sediment as varied with partition coefficients: A case study of polychlorinated dibenzodioxins/furans, polybrominated diphenyl ethers, phthalates, and polycyclic aromatic hydrocarbons. Sci. Total Environ. 2019, 670, 760–769.

53. Lesk, C.; Rowhani, P.; Ramankutty, N. Influence of extreme weather disasters on global crop production. Nature 2016, 529, 84.

54. Wang, D.; Felice, M. L.; Scow, K. M. Impacts and interactions of biochar and biosolids on agricultural soil microbial communities during dry and wet-dry cycles. Appl. Soil Ecol. 2020, 152, 103570.

55. Kim, S. W.; An, Y.-J. A simple and efficient method for separation of low-density polyethylene films into different micro-sized groups for laboratory investigation. Sci. Total Environ. 2019, 668, 84–89.

56. Brenner, S. J. The genetics of *Caenorhabditis elegans*. Genetics 1974, 77, 71–94.

57. Williams, P. L.; Dusenbery, D. B. Aquatic toxicity testing using the nematode, *Caenorhabditis elegans*. Environ. Toxicol. Chem. 1990, 9, 1285–1290.

58. Kim, S. W.; Moon, J.; An, Y.-J. A highly efficient nonchemical method for isolating live nematodes (*Caenorhabditis elegans*) from soil during toxicity assays. Environ. Toxicol. Chem. 2014, 34, 208–213.

59. Kim, S. W.; Moon, J.; An, Y.-J. Development of a nematode offspring counting assay for rapid and simple soil toxicity assessment. Environ. Pollut. 2018, 236, 91–99.

60. Reichardt, C. Solvents and Solvent Effects in Organic Chemistry. Wiley-VCH Publishers, 3rd ed., 2003.

61. International Organization for Standardization (ISO). Water quality—Determination of the toxic effect of sediment and soil samples on growth, fertility and reproduction of *Caenorhabditis elegans* (Nematoda). ISO 10872:2010. Geneva, Switzerland. 2010.

62. Acosta-Coley, I.; Duran-Izquierdo, M.; Rodriguez-Cavallo, E.; Mercado-Camargo, J.; Mendez-Cuadro, D.; Olivero-Verbel, J. Quantification of microplasatic along the Caribbean Coastline of Colombia: Pollution profile and biological effects on *Caenorhabditis elegans*. Mar. Pollut. Bull. 2019, 146, 574–583.

63. Kim, Y.; Jeong, J.; Lee, S.; Choi, I.; Choi, J. Identification of adverse outcome pathway related to high-density polyethylene microplastics exposure: *Caenorhabditis elegans* transcription factor RNAi screening and zebrafish study. J. Hazar. Mater, 2020, 388, 121725.

64. Schöpfe, L.; Menzel, R.; Schnepf, U.; Ruess, L.; Marhan, S.; Brümmer, F.; Pagel, H.; Kandeler, E. Microplastics effects on reproduction and body length of the soil-dwelling nematode Caenorhabditis elegans. Front. Env. Sci. 2020, 8, 41.

65. American Society for Testing Materials (ASTM). Standard guide for conducting laboratory soil toxicity tests with the nematode *Caenorhabditis elegans*. E2172–01. ASTM International, West Conshohocken, PA, USA. 2001.

66. Fueser, H.; Mueller, M.-T.; Weiss, L.; Höss, S.; Traunspurger, W. Ingestion of microplastics by nematodes depends on feeding strategy and buccal cavity size. Environ. Pollut. 2019, 255, 113227.

67. Mueller, M.-T.; Fueser, H.; Trac, L. N.; Mayer, P.; Traunspurger, W.; Höss, S. Surface-related toxicity of polystyrene beads to nematodes and the role of food availability. Environ. Sci. Technol. 2020, 54, 1790–1798.

68. Jang, M.; Shim, W. J.; Han, G. M.; Rani, M.; Song, Y. K.; Hong, S. H. Styrofoam Debris as a Source of Hazardous Additives for Marine Organisms. Environ. Sci. Technol. 2016, 50, 4951–4960.

69. Mercea, P. V.; Losher, C.; Herburger, M.; Piringer, O. G.; Tos, V.; Cassart, M.; Dawkins, G.; Faust, B. Repeated migration of additives from a polymeric article in food simulants. Polym. Test. 2020, 85, 106436.

70. IUPAC. Compendium of Chemical Terminology, 2nd ed. (the “Gold Book”). Compiled by A. D. McNaught and A. Wilkinson. Blackwell Scientific Publications, Oxford (1997). Online version (2019-) created by S. J. Chalk. ISBN 0-9678550-9-8. https://doi.org/10.1351/goldbook.

71. Zimmermann, L.; Dierkes, G.; Ternes, T. A.; Völker, C.; Wagner, M. Benchmarking the in Vitro Toxicity and Chemical Composition of Plastic Consumer Products. Environ. Sci. Technol. 2019, 53, 11467–11477.

72. Teuten, E. L.; Saquing, J. M.; Knappe, D. R. U.; Barlaz, M. A.; Jonsson, S.; Bjorn, A.; Rowland, S. J.; Thompson, R. C.; Galloway, T. S.; Yamashita, R.; Ochi, D.; Watanuki, Y.; Moore, C.; Viet, P. H.; Tana, T. S.; Prudente, M.; Boonyatumanond, R.; Zakaria, M. P.; Akkhavong, K.; Ogata, Y.; Hirai, H.; Iwasa, S.; Mizukawa, K.; Hagino, Y.; Imamura, A.; Saha, M.; Takada, H. Transport and release of chemicals from plastics to the environment and to wildlife. Phil. Trans. R. Soc. B 2009, 364, 2027–2045.

73. American Society for Testing Materials (ASTM). Standard Test Method for Leaching Solid Material in a Column Apparatus. D4874-95. ASTM International, West Conshohocken, PA, USA. 2014.

74. Organization for Economic Cooperation and Development (OECD). Guidelines for the testing of chemicals. Leaching in soil columns, 2004.

75. International Organization for Standardization (ISO). Soil quality — Leaching procedures for subsequent chemical and ecotoxicological testing of soil and soil-like materials — Part 1: Batch test using a liquid to solid ratio of 2 l/kg dry matter. ISO 21268-1:2019. Geneva, Switzerland. 2019a.

76. International Organization for Standardization (ISO). Soil quality — Leaching procedures for subsequent chemical and ecotoxicological testing of soil and soil-like materials — Part 2: Batch test using a liquid to solid ratio of 10 l/kg dry matter. ISO 21268-2:2019. Geneva, Switzerland. 2019b.

77. International Organization for Standardization (ISO). Soil quality — Leaching procedures for subsequent chemical and ecotoxicological testing of soil and soil-like materials — Part 3: Up-flow percolation test. ISO 21268-3:2019. Geneva, Switzerland. 2019c.

78. International Organization for Standardization (ISO). Soil quality — Leaching procedures for subsequent chemical and ecotoxicological testing of soil and soil-like materials — Part 4: Influence of pH on leaching with initial acid/base addition. ISO 21268-4:2019. Geneva, Switzerland. 2019d.

79. Galotto, M. J.; Torres, A.; Guarda, A.; Moraga, N.; Romero. Experimental and theoretical study of LDPE: Evaluation of different food simulants and temperatures. Food. Res. Int. 2011, 44, 3072–3078.

80. Li, B.; Wang, Z.-W.; Bai, Y.-H. Determination of the partition and diffusion coefficients of five chemical additives from polyethylene terephthalate material in contact with food simulants. Food Packag. Shelf life 2019, 21, 100332.

81. Mercea, P. V.; Kalisch, A.; Ulrich, M.; Benz, H.; Piringer, O. G.; Toşa, V.; Schuster, R.; Sejersen, P. Modelling migration of substances from polymers into drinking water. Part 2– Partition coefficient estimations. Polym. Test. 2019, 76, 420–432.

82. Tehrany, E. A.; Desobry, S. Partition coefficient of migrants in food simulants/polymers systems. Food Chem. 2007, 101, 1714–1718.

83. Endo, S.; Yuyama, M.; Takada, H. Desorption kinetics of hydrophobic organic contaminants from plastic pellets. Mar. Pollut. Bull. 2013, 74, 125–131.

84. Lee, H.; Byun, D.-E.; Kim, J. M.; Kwon, J.-H. Desorption modeling of hydrophobic organic chemicals from plastic sheets using experimentally determined diffusion coefficients in plastics. Mar. Pollut. Bullet. 2018, 126, 312–317.

85. Sun, B.; Hu, Y.; Cheng, H.; Tao, S. Releases of brominated flame retardants (BFRs) from microplastics in aqueous medium: Kinetics and molecular-size dependence of diffusion. Water Res.. 2019, 151, 215–225.

86. Tanaka, K.; Takada, H.; Yamashita, R.; Mizukawa, K.; Fukuwaka, M.; Watanuki, Y. Facilitated leaching of additive-derived PBDEs from plastic by seabirds’ stomach oil and accumulation in tissues. Environ. Sci. Technol. 2015, 49, 11799–11807.

