## Supplementary Material for "Effects of Different Microplastics on Nematodes in the Soil Environment: Tracking the Extractable Additives using an Ecotoxicological Approach"

6 pages

1 figure

2 tables

Reference list

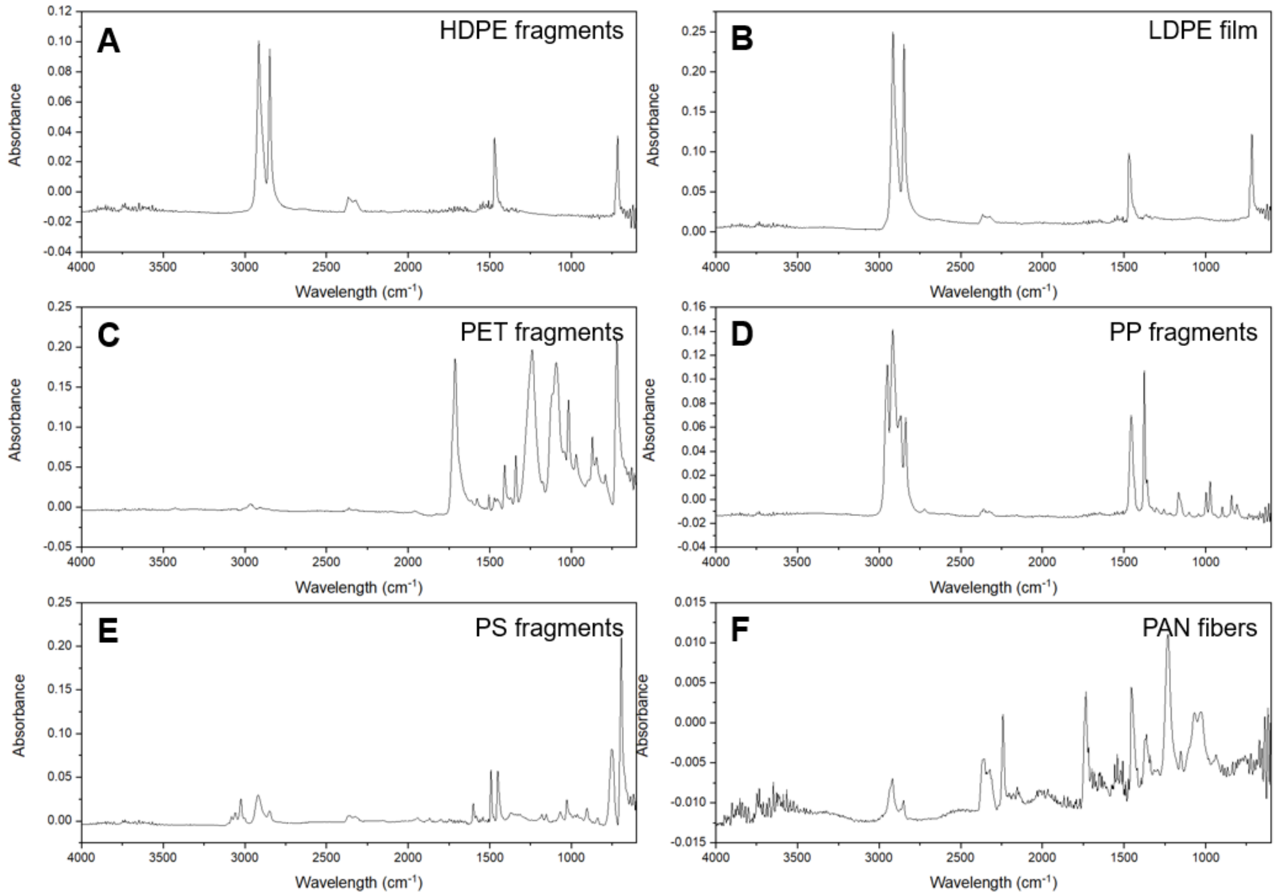


**Figure S1.** FTIR spectra, ATR mode, of all polymers tested in this paper: A) High-Density Polyethylene (HDPE); B) Low-Density Polyethylene (LDPE); C) Polyethylene Terephthalate (PET); D) Polypropylene (PP); E) Polystyrene (PS); F) Polyacrylonitrile (PAN).

**Table S1.** List of target plastic materials in this study.

| No. | Abbreviation | Composition | Shape | Size (μm) | Source | Ref. |
|---|---|---|---|---|---|---|
| 1 | HDPE | High-density polyethylene | Fragment (Irregularly) | 630-1000 | Bundesanstalt für Materialforschung und -prüfung (Berlin, Germany) | de Souza Machado et al. (2019) |
| 2 | | | | 250-630 | | |
| 3 | | | | <250 | | |
| 4 | PET | Polyethylene terephthalate | | 250-630 | | |
| 5 | | | | <250 | | |
| 6 | PP | Polypropylene | | 630-1000 | | |
| 7 | | | | 250-630 | | |
| 8 | | | | <250 | | |
| 9 | PS | Polystyrene | | 630-1000 | | |
| 10 | | | | 250-630 | | |
| 11 | | | | <250 | | |
| 12 | LDPE | Low-density polyethylene | Film | <630 | Commercial market (Seosan, South Korea) | Kim and An (2019) |
| 13 | PA | Polyacrylic | Fiber | <630 | Commercial market (Seoul, South Korea) | - |

**Table S2.** List of previous studies reporting microplastic toxicity on nematode *Caenorhabditis elegans*

| No. | Year | Author | Composition | Shape | Size | | Source | Test media | |
|---|---|---|---|---|---|---|---|---|---|
| 1 | 2017 | Zhao et al. | Polystyrene | Spherical | 108.2 | nm | Purchased | Liquid | K medium |
| 2 | 2018 | Dong et al. | Polystyrene | Particles | 108.2 | nm | Purchased | Liquid | Undefined |
| 3 | 2018a | Lei et al. | Polyamide<br>Polyethylene<br>Polypropylene<br>Polyvinyl chloride<br>Polystyrene<br>Polystyrene<br>Polystyrene | Fragment | <70<br>0.1<br>1<br>5 | µm | Undefined | Liquid | K medium |
| 4 | 2018b | Lei et al. | Polystyrene | Spherical | 100<br>500<br>1<br>2<br>5 | nm<br>µm | Purchased | Liquid | K medium |
| 5 | 2018 | Qu et al. | Polystyrene | Spherical | 108.2 | nm | Purchased | Liquid | K medium |
| 6 | 2019 | Acosta-Coley et al. | Polypropylene (predominant) | Pellet | 4.05-5.09 | mm | Collected | Liquid | K medium |
| 7 | 2019 | Fueser et al. | Polystyrene | Spherical | 0.5<br>1<br>3<br>6 | µm | Purchased | Liquid | K medium |
| 8 | 2019 | Kim et al. | Polystyrene | Spherical | 50<br>200 | nm | Purchased | Liquid | M9 buffer |
| 9 | 2019a | Qu et al. | Polystyrene | Spherical | 35 | nm | Synthesized | Liquid | K medium |
| 10 | 2019b | Qu et al. | Polystyrene | Spherical | 102.8 | nm | Purchased | Liquid | K medium |
| 11 | 2019c | Qu et al. | Polystyrene | Spherical | 101.6 | nm | Purchased | Liquid | K medium |
| 12 | 2019d | Qu et al. | Polystyrene | Spherical | 102.8 | nm | Purchased | Liquid | K medium |
| 13 | 2019e | Qu et al. | Polystyrene | Spherical | 102.8 | nm | Purchased | Liquid | K medium |
| 14 | 2020a | Kim et al. | Polystyrene | Spherical | 68<br>496 | nm | Purchased | Soil | LUFA |
| 15 | 2020b | Kim et al. | High-density polyethylene | Nonuniformly | 29-68 | µm | Purchased | Liquid | K media |
| 16 | 2020a | Li et al. | Polystyrene | Spherical | 32.45 | nm | Purchased | Liquid | K medium |
| 17 | 2020b | Li et al. | Polystyrene | Spherical | 98.7 | nm | Purchased | Liquid | K medium |
| 18 | 2020 | Mueller et al. | Polystyrene | Spherical | 0.1<br>0.5<br>1<br>3<br>6<br>10 | µm | Purchased | Liquid | M9 medium |
| 19 | 2020 | Qiu et al. | Polystyrene | Spherical | 102.8 | nm | Purchased | Liquid | K medium |
| 20 | 2020a | Qu et al. | Polystyrene | Spherical | 103.45 | nm | Purchased | Liquid | K medium |
| 21 | 2020b | Qu et al. | Polystyrene | Spherical | 103.45 | nm | Purchased | Liquid | K medium |
| 22 | 2020 | Schöpfer et al. | Low-density polyethylene<br>Polylactide/poly (butylene adipate-co-terephthalate) | Irregularly | 57<br>40 | µm | Purchased/milling | Liquid | M9 buffer |
| 23 | 2020 | Shang et al. | Polystyrene | Spherical | 1 | µm | Purchased | Agar | Nematode growth media |
| 23 | 2020 | Shang et al. | Polystyrene | Spherical | 5 | µm | Purchased | Agar | Nematode growth media |
| 24 | 2020 | Shao and Wang | Polystyrene | Spherical | 102.8 | nm | Purchased | Liquid | K medium |
| 25 | 2020 | Yang et al. | Polystyrene | Spherical | 102.8 | nm | Purchased | Liquid | K medium |
| 26 | 2020 | Yu et al. | Polystyrene | Spherical | 1.082 | µm | Purchased | Liquid | K medium |

**REFERENCES**

Acosta-Coley, I.; Duran-Izquierdo, M.; Rodriguez-Cavallo, E.; Mercado-Camargo, J.; Mendez-Cuadro, D.; Olivero-Verbel, J. Quantification of microplasatic along the Caribbean Coastline of Colombia: Pollution profile and biological effects on *Caenorhabditis elegans*. *Mar*. *Pollut*. *Bull*. **2019**, *146*, 574–583.

de Souza Machado, A. A.; Lau, C. W.; Kloas, W.; Bergmann, J.; Bachelier, J. B.; Faltin, E.; Becker, R.; Görlich, A. S.; Rillig, M. C. Microplastics can change soil properties and affect plant performance. *Environ*. *Sci*. *Technol*. **2019**, *53*, 6044−6052.

Dong, S.; Qu, M.; Rui, Q.; Wang, D. Combinational effect of titanium dioxide nanoparticles and nanopolystyrene particles at environmentally relevant concentrations on nematode *Caenorhabditis elegans*. *Ecotox*. *Environ*. *Safe*. **2018**, *161*, 444–450.

Fueser, H.; Mueller, M.-T.; Weiss, L.; Höss, S.; Traunspurger, W. Ingestion of microplastics by nematodes depends on feeding strategy and buccal cavity size. *Environ*. *Pollut*. **2019**, *255*, 113227.

Kim, S. W.; An, Y.-J. A simple and efficient method for separation of low-density polyethylene films into different micro-sized groups for laboratory investigation. *Sci*. *Total Environ*. **2019**, *668*, 84-89.

Kim, H. M.; Lee, D.-K.; Long, N. P.; Kwon, S. W.; Park, J. H. Uptake of nanopolystyrene particles induces distinct metabolic profiles and toxic effects in *Caenorhabditis elegans*. *Environ*. *Pollut*. **2019**, *246*, 578−586.

Kim, S. W.; Kim, D.; Jeong, S.-W.; An, Y.-J. Size-dependent effects of polystyrene plastic particles on the nematode *Caenorhabditis elegans* as related to soil physicochemical properties. *Environ*. *Pollut*. **2020a**, *258*, 113740.

Kim, Y.; Jeong, J.; Lee, S.; Choi, I.; Choi, J. Identification of adverse outcome pathway related to high-density polyethylene microplastics exposure: *Caenorhabditis elegans* transcription factor RNAi screening and zebrafish study. *J. Hazar*. *Mater*, **2020b**, *388*, 121725.

Lei, L.; Wu, S.; Lu, S.; Liu, M.; Song, Y.; Fu, Z.; Shi, H.; Raley-Susman, K. M.; He, D. Microplastic particles cause intestinal damage and other adverse effects in zebrafish *Danio rerio* and nematode *Caenorhabditis elegans*. *Sci*. *Total Environ*. **2018a**, *619–620*, 1–8.

Lei, L.; Liu, M.; Song, Y.; Lu, S.; Hu, J.; Cao, C.; Xie, B.; Shi, H.; He, D. Polystyrene (nano)microplastics cause size-dependent neurotoxicity, oxidative damage and other adverse effects in *Caenorhabditis elegans*. *Environ*. *Sci*. *Nano* **2018b**, *5*, 2009.

Li, D.; Deng, Y.; Wang, S.; Du, H.; Xiao, G.; Wang, D. Assessment of nanopolystyrene toxicity under fungal infection condition in *Caenorhabditis elegans*. *Ecotoxicol*. *Environ*. *Safe*. **2020a**, *197*, 110625.

Li, D.; Ji, J.; Yuan, Y.; Wang, D. Toxicity comparison of nanopolystyrene with three metal oxide nanoparticles in nematode *Caenorhabditis elegans*. *Chemosphere* **2020b**, *245*, 125625.

Mueller, M.-T.; Fueser, H.; Trac, L. N.; Mayer, P.; Traunspurger, W.; Höss, S. Surface-related toxicity of polystyrene beads to nematodes and the role of food availability. *Environ*. *Sci*. *Technol*. **2020**, *54*, 1790-1798.

Qiu, Y.; Luo, L.; Yang, Y.; Kong, Y, Li, Y.; Wang, D. Potential toxicity of nanopolystyrene on lifespan and aging process of nematode *Caenorhabditis elegans*. *Sci*. *Total Environ*. **2020**, *705*, 135918.

Qu, M.; Xu, K.; Li, Y.; Wong, G.; Wang, D. Using *acs*-22 mutant *Caenorhabditis elegans* to detect the toxicity of nanopolystyrene particles. *Sci*. *Total Environ*. **2018**, *643*, 119-126.

Qu, M.; Qiu, Y.; Kong, Y.; Wang, D. Amino modification enhances reproductive toxicity of nanopolystyrene on gonad development and reproductive capacity in nematode *Caenorhabditis elegans*. *Environ*. *Pollut*. **2019a**, *254*, 112978.

Qu, M.; Zhao, Y.; Zhao, Y.; Rui, Q.; Kong, Y.; Wang, D. Identification of long non-coding RNAs in response to nanopolystyrene in *Caenorhabditis elegans* after long-term and lowdose exposure. *Environ*. *Pollut*. **2019b**, *255*, 113137.

Qu, M.; Nida, A.; Kong, Y.; Du, H.; Xiao, G.; Wang, D. Nanopolystyrene at predicted environmental concentration enhances microcystin-LR toxicity by inducing intestinal damage in *Caenorhabditis elegans*. *Ecotox*. *Environ*. *Safe*. **2019c**, *183*, 109568.

Qu, M.; Luo, L.; Yang, Y.; Kong, Y.; Wang, D. Nanopolystyrene-induced microRNAs response in *Caenorhabditis elegans* after long-term and lose-dose exposure. *Sci*. *Total Environ*. **2019d**, *697*, 134131.

Qu, M.; Kong, Y.; Yuan, Y.; Wang, D. Neuronal damage induced by nanopolystyrene particles in nematode *Caenorhabditis elegans*. *Environ*. *Sci*. *Nano* **2019e**, *6*, 2591.

Qu, M.; Li, D.; Zhao, Y.; Yuan, Y.; Wang, D. Exposure to low-dose nanopolystyrene induces the response of neuronal JNK MAPK signaling pathway in nematode *Caenorhabditis elegans*. *Environ*. *Sci*. *Eur*. **2020a**, *32*, 58.

Qu, M.; Li, D.; Qiu, Y.; Wang, D. Neuronal ERK MAPK signaling in response to low-dose nanopolystyrene exposure by suppressing insulin peptide expression in *Caenorhabditis elegans*. *Sci*. *Total Environ*. **2020b**, *724*, 138378.

Schang, X.; Lu, J.; Feng, C.; Ying, Y.; He, Y.; Fang, S.; Lin, Y.; Dahlgren, R.; Ju, J. Microplastic (1 and 5 μm) exposure disturbs lifespan and intestine function in the nematode *Caenorhabditis elegans*. *Sci*. *Total Environ*. **2020**, *705*, 135837.

Schöpfe, L.; Menzel, R.; Schnepf, U.; Ruess, L.; Marhan, S.; Brümmer, F.; Pagel, H.; Kandeler, E. Microplastics effects on reproduction and body length of the soil-dwelling nematode Caenorhabditis elegans. *Front*. *Env*. *Sci*. **2020**, *8*, 41.

Shao, H.; Wang, D. Long-term and low-dose exposure to nanopolystyrene induces a protective strategy to maintain functional state of intestine barrier in nematode *Caenorhabditis elegans*. *Environ*. *Pollut*. **2020**, *258*, 113649.

Yang, Y.; Shao, H.; Wu, Q.; Wang, D. Lipid metabolic response to polystyrene particles in nematode *Caenorhabditis elegans*. *Environ*. *Pollut*. **2020**, *256*, 113439.

Yu, Y.; Chen, H.; Hua, X.; Dang, Y.; Han, Y.; Yu, Z.; Chen, X.; Ding, P.; Li, H. Polystyrene microplastics (PS-MPs) toxicity induced oxidative stress and intestinal injury in nematode *Caenorhabditis elegans*. *Sci*. *Total Environ*. **2020**, *726*, 138679.

Zhao, L.; Qu, M.; Wong, G.; Wang, D. Transgenerational toxicity of nanopolystyrene particles in the range of μg $L^{-1}$ in the nematode *Caenorhabditis elegans*. *Environ*. *Sci*. *Nano* **2017**, *4*, 2356.
